# Probing the Biology of Zinc Alpha2-Glycoprotein and the role it plays in cachexia

**DOI:** 10.1101/2022.12.14.520465

**Authors:** Parth Pandit, Subrat Panigrahi

## Abstract

Cachexia is a metabolic disease that results in drastic weight loss and muscle wasting. 20% of total cancer patients will die due to cachexia related complications. ZAG contributes to the regulation of weight and body fat through lipid and glucose metabolism. In healthy individuals, ZAG exerts a homeostasis effect by inducing lipolysis of adipose tissue to help reduce fat storage and overall weight. ZAG is upregulated in various carcinomas and cancer patients with upregulated ZAG are observed to lose weight rapidly. The mutants of ZAG which are the 4 amino acids Tryptophan 148, Arginine 73, Phenylalanine 101, Isoleucine 76 have all been mutated to Alanine. The effect of mutants and the wild type ZAG can also be found out by conducting experiments. ZAG has a potential lipid binding site that could be imperative to the function of ZAG. A lipolysis colorimetric kit allows us to measure the ZAG variants effects on 3T3 adipose cells to determine what β-adrenoreceptor signaling pathways are being utilized in its lipolytic effect. The Tryptophan-Alanine mutant showed increased lipolysis after 1h treatment than other variants. The time period is also a rate limiting step which can play major factor in lipolysis. The ultimate goal is to identify the ligand(s) and the interactions between them and ZAG. The design of a therapeutic would give patients options of treatments brought about by attenuating the weight loss. With this, it would offer a better prognosis for patients and provide them with a greater quality of life.

## Introduction

Cancer accounts for one of the major causes of death worldwide which accounts for nearly 10 million deaths in 2020. Around one-third of deaths from cancer are due to consumption of tobacco and alcohol, high body mass index, low appetite and satiety, and lack of exercise[1]. There are many symptoms which can be observed at different stages of cancer such as pain or ache, unexplained weight loss, swelling and fatigue [2].

### Cachexia

An unintentional weight loss which can be prominently seen in any chronic disease is referred to as wasting syndrome or Cachexia [3], Weight loss can arise from a variety of reasons, but cachexia affects you regardless of how much you eat. Cachexia causes both fat and muscle loss. The fact that cachexia occurs unintentionally and distinguishes it from typical weight loss. Simultaneously, metabolism is altered leading to break down of fats in the body. The appetite is also affected by inflammation, and the body burns more calories than it should. Cachexia accounts for nearly 80% cancer patients of which 20% ultimately leads to death in cancer patients. It can also be seen in chronic obstructive pulmonary disease and chronic kidney disease. Cachexia prevalence increases with the progress of cancer in patients [4]. The consequences of the wasting disorder are reduction of response to therapy and treatment tolerance, reduced quality of life with disability and shortened survival. Cachexia is dependent on multiple factors such as abnormal metabolism induced by tumor and host derived factors and inflammation which causes adipose wastage.

### Role of Adipose tissue

Adipose tissue plays a vital role in tumour microenvironment. These are large, interactive fat depot sites. White Adipose Tissue (WAT) comprises of largest section of the tissue and is regarded as energy storage in the form of triglycerides. WAT plays both endocrine as well as paracrine role. WAT is found in visceral, subcutaneous regions. On the other hand, Brown Adipose tissue (BAT) functions to spend energy. BAT deposits are found around the aorta and in the neck’s supraclavicular area [5]. The presence and activation of a proton leakage route mediated by uncoupling protein 1 (UCP1) - the biomarker of BAT function - is largely responsible for the presence and ability of these adipocytes to spend substantial amounts of energy. UCP1 decouples oxidative phosphorylation from ATP production in the inner mitochondrial membrane, allowing heat to be transferred. The new beige adipocytes are classical drivers for cachexia in tumour microenvironment. These beige adipocytes are derived from precursors of adipose [6] **(figure1**). They function same as BAT as energy spenders but share a similar morphology with WAT. The Early B cell factor 2 (Ebf2) along with Peroxisome proliferator–activated receptor γ (PPARγ) help in induction of PR domain containing 16 (Prdm16) protein. This domain along with Ebf2 induces adipocytes differentiation and causes browning of white adipose cells [7]. The browned adipocytes eat up the glucose and fatty acids and dissipate energy in the form of heat. The secretions of adipose tissue are mainly a wide array of proteins called as adipokines [8]. These adipokines such as leptin, adiponectin, TNF-α, visfatin, ZAG have a mutual connection with brain, liver, muscle to influence appetite, food metabolism, insulin sensitivity, inflammation, and the formation of adipose tissue [9, 10].

**Figure 1.**
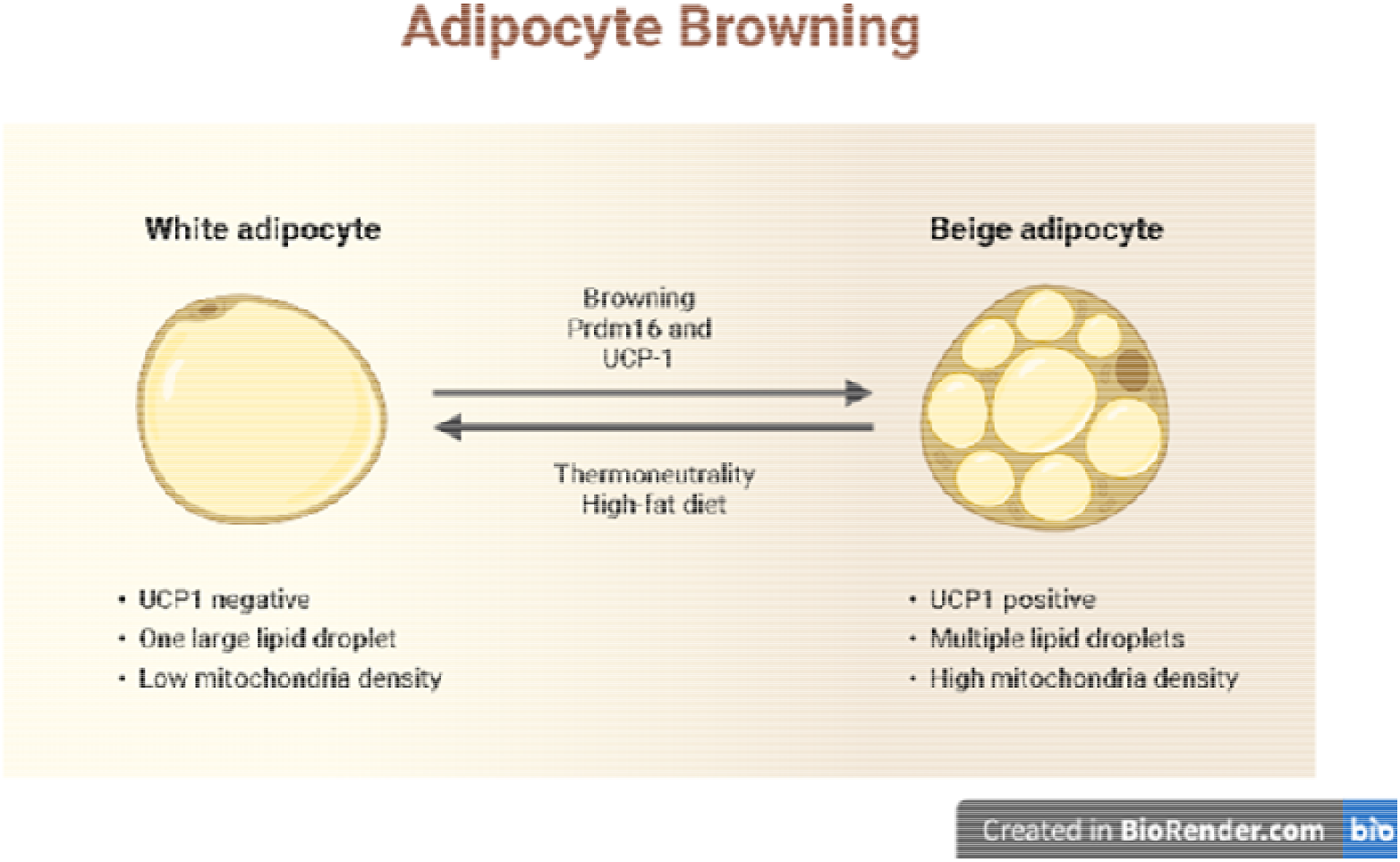
Adipocyte browning with the help of Prdm 16 - PR/SET domain containing 16 and UCP-1 - Uncoupling protein

### Zinc Alpha 2 Glycoprotein

Along with many adipokines which play a role in cachexia such as TNFα, IL-1 β and IL-6. ZAG is 43 kDa molecular protein found in human plasma and encoded by AZGP1 gene with 295-amino acid-long polypeptide **(figure2**). The crystal structure of ZAG bears a resemblance to that of Major Histocompatibility Class I protein (MHC) which has grooves of α1, α2 and α3 and two β sheet domains. The ligand binds to α1, α2 only by hydrophobic linkages. Sequence homology shared is 30% to 47% similar to classic MHC-I like molecules, which play an essential role in antigen presentation and processing by cytotoxic T cells. Lipid-Mobilizing Factor is a protein that is secreted from the cancer cells and has the ability for causing wastage of adipose. ZAG is produced by epithelial cells of the liver and prostate in both in WAT and BAT .ZAG is secreted from these cells and its expression has been reported in blood and other body fluids such as cerebrospinal fluid, milk and sweat [11]. The Zinc-α2-Glycoprotein (ZAG) which resembles to Lipid-Mobilizing Factor (LMF) in the amino acid sequence and immunoreactivity is vital for cachexia [12]. The concentration of ZAG increases with age which is highest in adults and lowest in fetal stage. The concentration is seen to increase in tumour of different organs hence it is a biomarker for carcinoma. Hence, upregulation of this glycoprotein results in increased weight loss in cancer. Silencing ZAG results in termination of lipolysis. ZAG stimulates adipose wastage by adipocytes both in vivo and in vitro. Lipolysis is controlled by c-AMP dependent protein kinase A’s reversible phosphorylation of a single serine residue. The concentration of c-AMP in adipocytes is proportional to adipocyte fat loss [13].

**Figure 2.**
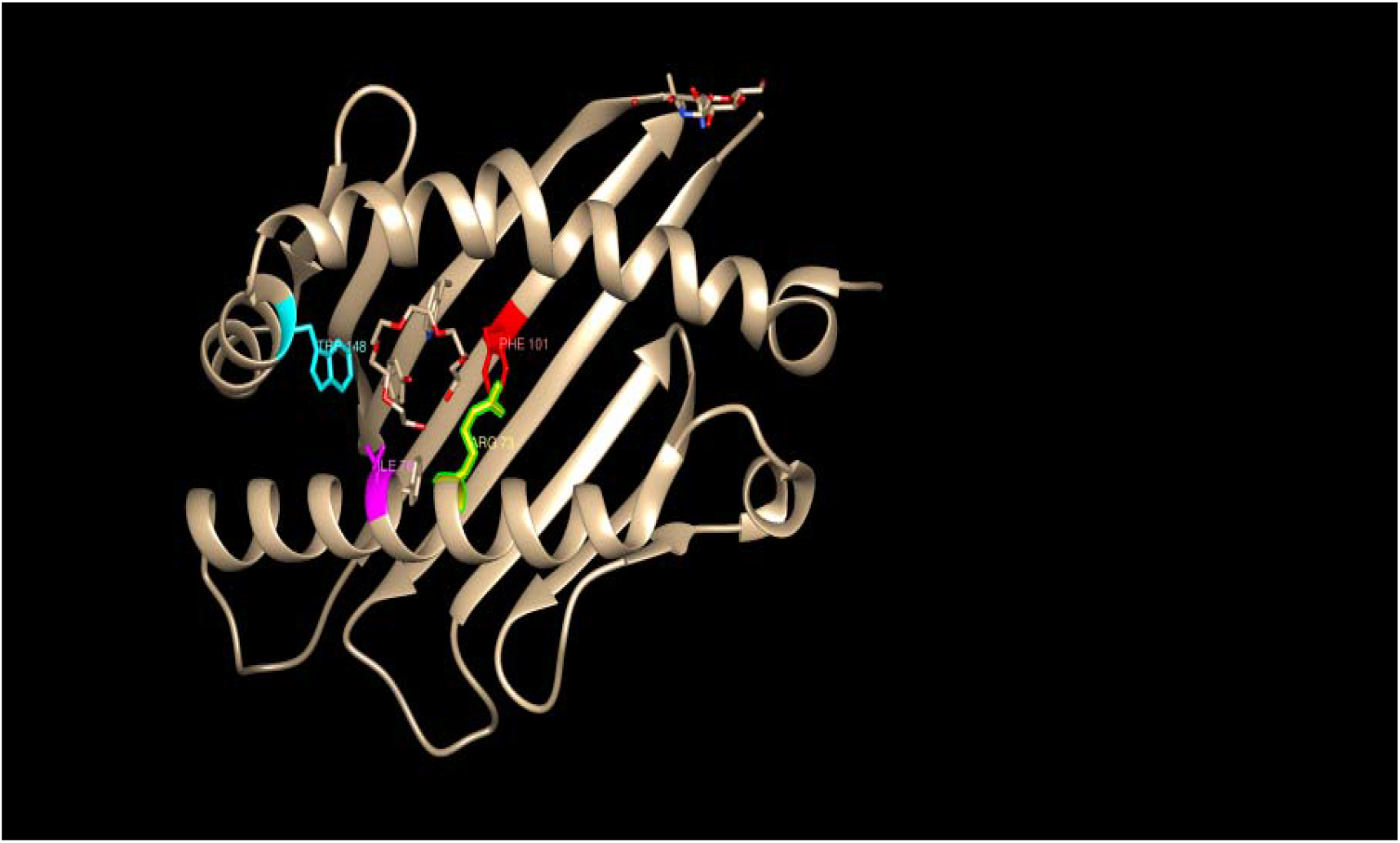
Zinc Alpha 2 Glycoprotein crystal structure obtained from protein data bank (1T7V) showing the four amino acid residues essential for binding to any fatty acid ligand Blue- Tryptophan, Red -Phenylalanine, Yellow- Arginine, Magenta- Isoleucine.

### Effect of ZAG on adiposity

ZAG works by the mechanism of browning of WAT (figure1). This browning takes place by stimulation of PPAR γ and early B cell factor 2 which in turn increase the binding to PRDM16 and UCP1. These Prdm16 and UCP1 are known to convert white adipocytes into brown-like adipocytes and promote energy consumption. Mechanism is mediated by Protein Kinase A (PKA) and p38 mitogen-activated protein kinase (MAPK) signaling, which upregulates ZAG expression of many lipolysis causing molecules [14]. ZAG also helps in growth of 3T3-L1 preadipocytes but inhibits their differentiation. The pharmacological action of loss of adipose and with oxygen consumption in BAT along with increased glycerol levels is due to β-3 adrenergic receptor. Lipolysis is exerted through Cyclic AMP pathway through β-3 adrenergic receptor. LMF also works by the mechanism of activation of lipolysis in adipocytes by GTP dependent adenylate cyclase process [15]. Expression of ZAG causes lot of changes such as enhanced lipolysis, decreased lipogenesis, and decreased lipoprotein lipase (LPL) have all been linked to loss of triglycerides, which would inhibit synthesis of lipid. Glycerol and free fatty acids (FFA) turnover are high, and an increase in mobilization of lipids in cancer patients is evident. Increased production and activity of hormone-sensitive lipase (HSL), a rate-limiting enzyme in the lipolytic pathway, is predicted to accelerate lipolysis. Lipolysis will see an increase in hormone-sensitive lipase mRNA and protein levels. Increased production and activity of HSL, a rate-limiting enzyme in the lipolytic pathway, is predicted to accelerate lipolysis[10]. Lipolysis will see an increase in hormone-sensitive lipase mRNA and protein levels. Fatty Acid Synthase (FAS), acetyl-CoA carboxylase, and acyl-coenzyme A, are also downregulated with ZAG expression. There is a distinctive browning of adipose tissue which in turn leads to energy expenditure and heat production in cachexia caused by cancer [9, 16]. Overall, the body temperature increases and homeostatic imbalance is observed with decrease in body weight by expression of ZAG. ZAG binds to new binding entity called Amine Oxidase Copper-containing 3 (AOC3). This enzyme is located in the cell surface of adipocytes and inhibition of this enzyme by ZAG leads to fibrosis [17].

### ZAG structural biology

This Glycoprotein has enfolded the zinc ion which is predicted to bind close to its groove. ZAG has 15 weak binding sites which are responsible for separation from plasma. ZAG is soluble and in its α-helical groove, it prefers tiny hydrophobic ligands to antigenic peptides. It has 4 amino acid residues to which ligand binds to are Arginine (Arg), Tryptophan (Trp), Isoleucine (Ile), Phenylalanine (Phe). The crystallized structure of ZAG is obtained from the protein data bank, (PDB ID-1T7V). The exogenous ligand which is bound in pocket with hydrophobic linkage is hexaethylene glycol (P6G). More crystal work was analyzed by fluorescent ligand (DAUDA) 11-(dansylamino) undecanoic acid in the grove between the α1 and α2 domains. There is tetramer formation in the presence of DAUDA [18]. The ligand C16-BODIPY which is fluorescent labelled also bind to ZAG[19]. Thus, there are notable aims and hypothesis needs to be proven to understand the biology and binding studies of ZAG.

## Materials and Methods

### Bacterial transformation

BL21 E. coli cells were incubated with pQE60 plasmid designed to express his-tagged ZAG and were inoculated for 5 min in an Eppendorf tube on ice. Media used in this experiment is 2YT which is composed of 1.6% Tryptone (w/v), 1% Yeast extract (w/v) and 0.5% Sodium Chloride (NaCl) (w/v) in distilled water. After incubation samples were heat-shocked at 42°C for 45 sec then returned to ice for 2 mins. Media was added and after a 1h incubation the mixture was spread evenly on Amp plates. The spreader was passed through the Bunsen after each plate to prevent cross contamination. Plates were then covered in parafilm and incubated at 37°C overnight for growth of single colonies. Colony growth results from a successful transformation as the pQE60 plasmid expresses ampicillin resistance.

### Miniprep DNA purification method

Single colonies were taken from the transformed plate and the pipette is dipped in 20mL falcon tubes containing 2YT media and ampicillin. The tubes were kept overnight in shaking incubator at 37°C. These cultures underwent Wizard Plus SV Minipreps DNA purification System (Promega, #TB225). This is carried out by production of clear lysate. The pellet was resuspended and mixed thoroughly in 250μL in Cell resuspension solution along with 250μL of cell lysis solution. Alkaline protease solution of 10μL is added to the mixture and incubated 5 mins. 350μL of Neutralization solution is added and centrifuged at top speed for 10 mins and transferred to Eppendorf tubes. A spin column was placed in a collecting tube, and the cell lysate was deposited into the spin column and centrifuged at top speed for 1 minute at room temperature in a microfuge. The flow through was removed. Add 750μL of wash solution and centrifuge at high speed for 1 min before repeating with 250μL of wash solution This solution is allowed to centrifuge at top speed for 2 min. The solution is transferred to the microcentrifuge tube (1.5mL) and 100μL Nuclease free water is added to it. After centrifugation for 1 min at top speed, the solution which is eluted is DNA plasmid pQE60 which is stored at −20°C.

### Restriction Digest Analysis

Gel preparation for restriction digest analysis is made out of Agarose gel (0.8% agarose powder (w/v), Trichloroacetic acid (TAE buffer)). This gel is microwaved and stirred at regular intervals till it becomes transparent liquid. The liquid is allowed to cool and poured into the gel casket. Ethidium bromide 1.5μL is added to it and mixed until a clear gel is not obtained. A comb was inserted and left until it was set. 5μL of 1 kb DNA ladder and 10μL of each sample were pipetted into the wells, and the gel was run for 30 min at 75 volts. After running the samples, the gel was placed under U.V light to image the bands. Single as well as double digests were carried out from the miniprep samples. The enzymes included were Bam HI, Eco RI, Bgl III and Hind III. The simulated version of this was made up using Snapgene software (version 6.2).

### Protein Purification

Transformed BL21-PQE60 ZAG expressing single colonies were taken from plate using a pipette tip and dispensed into 20 mL falcon tubes with 2YT media and ampicillin (100μg/mL). The resulting culture was then transferred to conical flask containing 400mL of 2YT media and ampicillin grown overnight. The culture was transferred to 500mL drums and spun 3500xg for 20 min at room temperature in worktop centrifuge. This process gives out a resultant pellet. The pellet was combined with lysis solution (50 mM Tris-HCl, pH 8.0, 25% sucrose (w/v), 1 mM EDTA), and cells were lysed with lysozyme (1 mg) dissolved in lysis buffer before being incubated on ice for 30 min. The presence of viscosity is due to the inclusion of lysozyme. The mixture was then incubated on ice for 30 min with MgCl2 (10 mM), MnCl2 (1 mM), and DNase I (10 μg/mL). Viscosity of the solution decreases as DNase I is added. Detergent buffer (0.2 M NaCl, 1% deoxycholic acid (w/v), 1% Nonidet P-40 (v/v), 20 mM Tris-HCl (pH 7.5), 2 mM EDTA) was added to the cell lysate, incubated for 20 min and then centrifuged at top speed for 10 min at 4°C. The supernatant is discarded and five washes of 0.5% Triton X-100, 1 mM EDTA solution. This process after repeating continuously gives us a tighter pellet.

The samples after washing were transferred to Eppendorf tubes of 1.5mL and then can be spun at top speed for 10 mins at 4°C. This is done in order to bring down all the protein into the pellet. The denaturation of the protein is done by 1mL of 8M urea or 1mL of Guanidium Hydrochloride to remove recombinant protein from inclusion bodies and 3 mL of Refolding buffer (0.1 M Tris-HCl, 2mM EDTA, 0.4 M L-arginine, 0.5 mM oxidized glutathione, 5 mM reduced glutathione) was added and incubated overnight. The first was taking 1mL of the overnight incubation and placing it in vivaspin 20 centrifugal concentrator. Refolding buffer was gradually added to a total of 20 mL. The sample was then spun at 3500xg until 5 mL of concentrated ZAG solution was left in the top part of the vivaspin tube. This sample was then collected and analyzed for protein concentration in Nanodrop (ThermoFisher, 701-058112). The proteins were measured at wavelength of 280nm UV visible spectrum and which enables high speed accuracy of 1.5μL samples. The concentrations are measured in mg/mL. The integral protein structure was made up by using UCSF chimera (version 1.16).

### Lipolysis assay

It is a colorimetric assay kit (Sigma Aldrich MAK211) that contains synthetic catecholamine isoproterenol that stimulates β adrenergic receptors. This 3T3-L1 Adipocytes were assessed in this assay. The 3T3-L1 preadipocytes are mainly fibroblasts which are grown and differentiated in 12-well cell culture plates in presence of media Fetal Bovine Serum (FBS) [20]. Day-10 cells after adipogenic induction were used for the lipolysis assay. After the differentiation, wash the cells two times with 100μL of lipolysis assay buffer. Remove the wash buffer and replace it with 150μL Lipolysis Assay Buffer. Addition of 1.5μL of 10μM isoproterenol which stimulates lipolysis. 20μL of media (cells) is being added into 96 well plate with addition of 30μL of lipolysis assay buffer so that the total reaches to 50μL. The reaction mix of glycerol is being made up by making the total to 50μL with fixed mixture of 46μL of glycerol assay buffer and 2μL each of glycerol probe and glycerol enzyme mix to the 96 well titre plate. The plate is incubated at room temperature for 30 mins protected from light. The absorbance is read at 570nm in a microtiter plate reader.

### Oil Red O staining

Oil red O staining is used to quantify neutral lipid droplets (Sigma Aldrich MAK194). 3T3 L1 adipocytes grown and differentiated on 12 well plates were used, with triplicate determinations for each condition (no ZAG, ZAG, mutant ZAG etc.). After ZAG treatment as indicated on the figures, the media was aspirated and cells were incubated in 10% (v/v) formalin. Cells were washed in 60% isopropanol and then Oil Red O (0.2%) v/v added for 10 minutes. Cells were then washed with water 4 times. 1mL of isopropanol (VWR 67-63-0) was used to elute the stain. 100μL aliquots were added to a 96-well plate and absorbance measured at 540nm. The 12-well cell plate was also imaged with EVOS® FL Auto Imaging System (Leica) to image the stained lipid droplets at 10x or 20x magnification.

### Protein analysis by western blotting

The stand for gel to be placed with glass slides of 1.5mm for the gel to be set. The running gel and stacking gel is made with given ingredients. Tris-glycine SDS-polyacrylamide running gel (1.5M Tris-HCl, pH8.8, 0.1% (v/v), 10% SDS, 30% (v/v) acrylamide mix, 10% (v/v) ammonium persulfate and 0.1% (v/v) TEMED) (Thermofisher,17919) combined with 5% stacking gel; (1.0M Tris-HCl, pH6.8, 0.1% (v/v), 10% SDS, 30% (v/v) acrylamide mix, 10% (v/v) ammonium persulfate and 0.1% (v/v) TEMED). Gels were run in SDS-Running buffer (0.19M glycine, 25mM Tris-HCl, and 0.1% (v/v) SDS) at 150V for 2-h period. Running buffer is put with TEMED in it and brought to level till the top with water. After removing the water, stacking buffer is added with TEMED and wells are inserted quickly. The gels are left untouched till it solidifies and wells are formed. 3μL of pre-stained protein molecular weight Precision Plus Protein™ Standard ladder (Bio-Rad, 1610374) and 10μL of each sample is added to 15 well polyacrylamide gels 10% (w/v) using a 20 μL pipette. Gels were secured in clamp stands and submerged with running buffer comprising of glycine (0.19 M), SDS 0.1% (w/v) and Trizma®base (25 mM). The gel apparatus was then attached to a power source and was set at 100 V for 60 min and then increased to 150 V when samples had run halfway down the gel.

### Wet transfer and immunoblotting

Once the separation was complete, the gel was placed into a transfer cassette against nitrocellulose paper (ThermoFisher, LC2006) via the use of Mini Trans-Blot® Cell apparatus (Bio-Rad Laboratories, California) following manufacturers protocol, supported with sponge and card. The assembled transfer cassette was submerged in transfer buffer consisting of glycine (0.19 M), Trizma®base (25 mM) and methanol 20% (v/v). Protein transfer from the gel to the nitrocellulose was at 60 V for 135 min. The transferred blot is then poured with ponceau dye (0.1% (v/v) Ponceau S, 5% (v/v) acetic acid) (ThermoFisher, A40000279) to visualize the proteins run on the paper. The dye is then washed with PBS. The nitrocellulose paper was incubated in blocking buffer comprising of Bovine Serum Albumin (BSA) 3% (w/v) in TBST (20mM Tris-HCl, 150mM NaCl, 0.1% Tween (v/v) for 1 h. The membrane was then placed in appropriate primary antibody (1:1000) diluted in BSA 1% (w/v), TBST and rotated overnight at 4°C. After primary incubation, blots underwent three 5-minute washes with TBST or PBST whilst rocking. They were then incubated with the corresponding LI-COR IR Dye-conjugated anti-mouse/rabbit secondary antibody (1:10,000) diluted in BSA 3% (w/v) and TBST for 1 h at room temperature (23°C). The antibodies used were mostly rabbit as it gives us a bright green fluorescence. Subsequently, blots underwent another three 5-minute wash steps in TBST. Proteins were visualized using the LI-COR Infrared 9120 imaging system (Odeyssesy®) at 700nm and 800 nm channels following manufacturers protocol. Data was analyzed via Image Studio Lite (Cambridge, UK) imaging system.

## Results

### Crystal structure of ZAG protein

The protein structure of ZAG was retrieved from the protein data bank (PDB ID-1T7V) and all visualizations performed in UCSF Chimera. As described, ZAG is a 43 kDa soluble protein homologous to the MHC class I heavy chain. ZAG can bind to small hydrophobic ligands [21] although the biological ligand remains to be identified. The fluorophore-tagged fatty acid ligand DAUDA exhibits a substantial increase in fluorescence and blue-shift is observed upon binding to ZAG. The X-ray crystal structure solved for ZAG displays a polyethylene glycol molecule (P6G) which is likely substituting for a higher affinity natural ligand, occupying an apolar groove between its α domain helices, which matches to the peptide binding groove in class I MHC proteins. ZAG binds differently to boron dipyrromethene C-16 BODIPY. In the presence of increasing concentration of zinc, ZAG binding to C16-BODIPY is reduced compared to DAUDA [22]. We conclude that the lipid-binding groove in ZAG contains at least two distinct fatty acid-binding sites for DAUDA and C16-BODIPY, similar to the multiple lipid binding seen in the structurally related immune protein CD1c. The fluorophore displacement shows that ZAG needs long chained fatty acid ligand which can bind in the pocket. The four amino acid residues to which every ligand bind is Arginine 73, Tryptophan 148, Phenylalanine 101, Isoleucine 76. The deliberate mutation of these residues to alanine may or may not disrupt the ligand binding and in turn has that resulting effect on lipolysis. This hypothesis can be more understood by further investigating into the ligand and protein studies and its effect on the adipose cells (**figure 2,3,4,5,6**).

**Figure 3.**
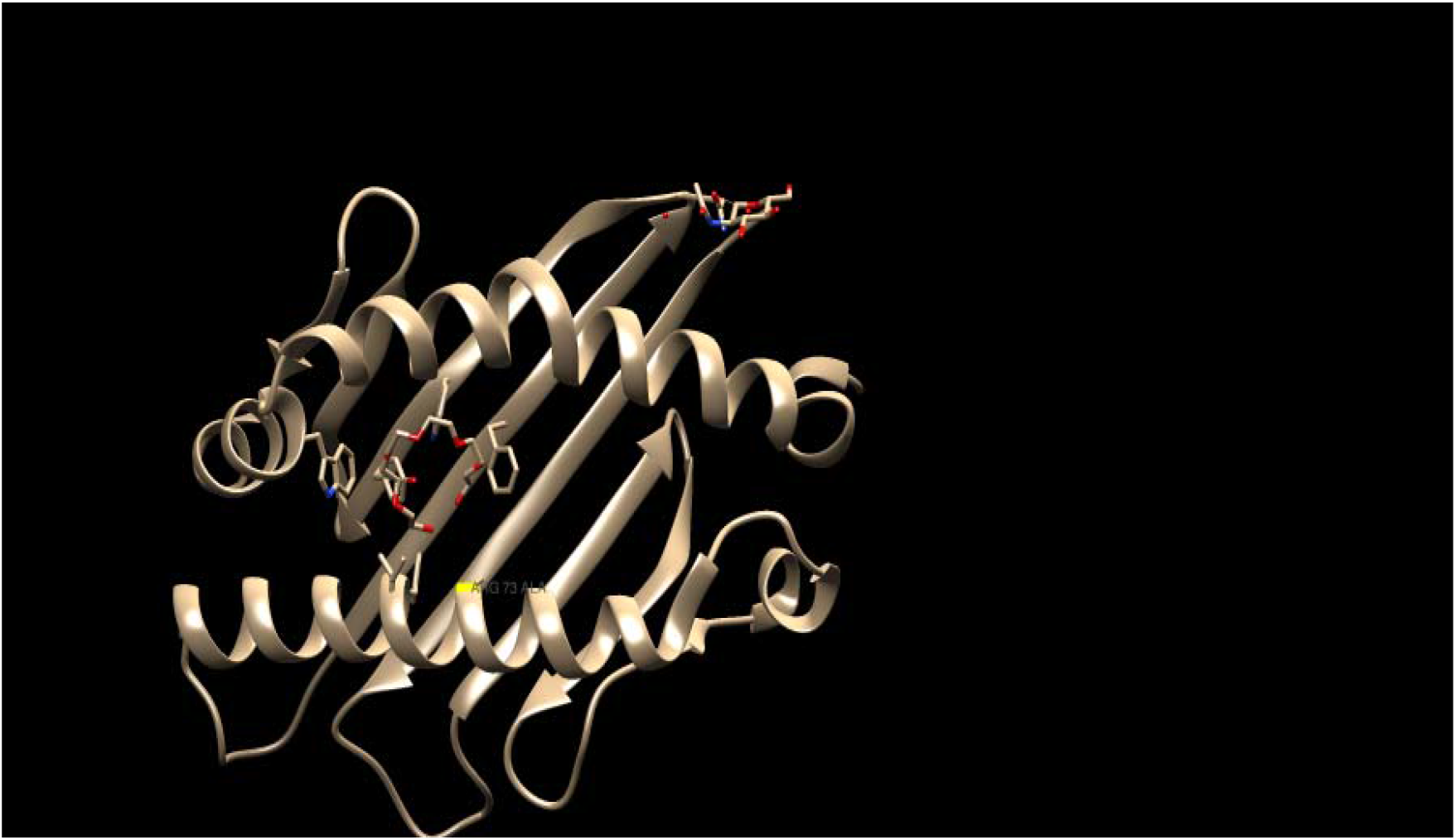
Zinc Alpha 2 Glycoprotein crystal structure obtained from protein data bank (1T7V) showing the Arginine amino acid residue mutated to Alanine. Blue- Tryptophan, Red -Phenylalanine, Yellow- Arginine, Magenta- Isoleucine.

**Figure 4.**
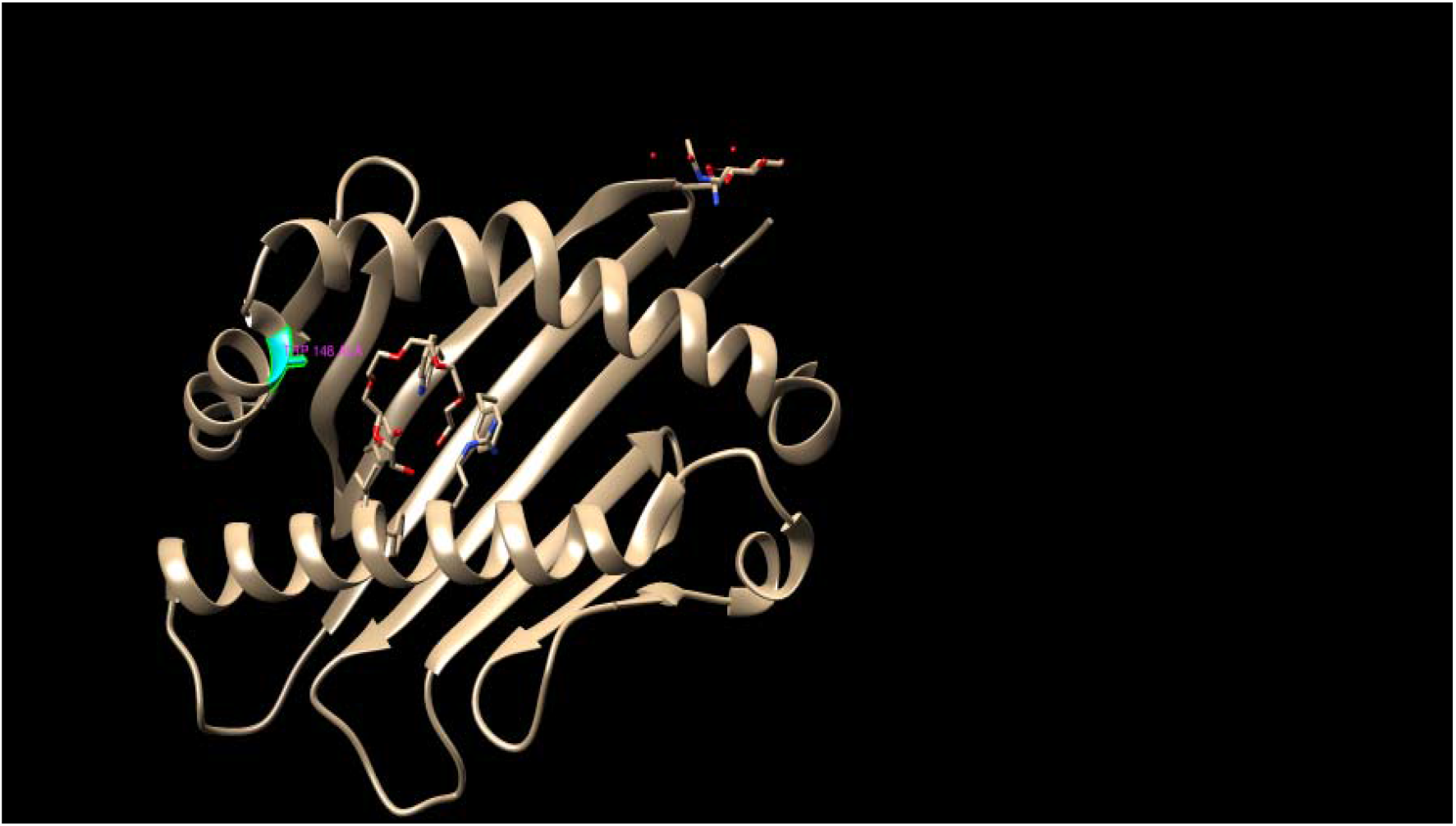
Zinc Alpha 2 Glycoprotein crystal structure obtained from protein data bank (1T7V) showing the Tryptophan amino acid residue mutated to Alanine. Blue- Tryptophan, Red -Phenylalanine, Yellow- Arginine, Magenta- Isoleucine.

**Figure 5.**
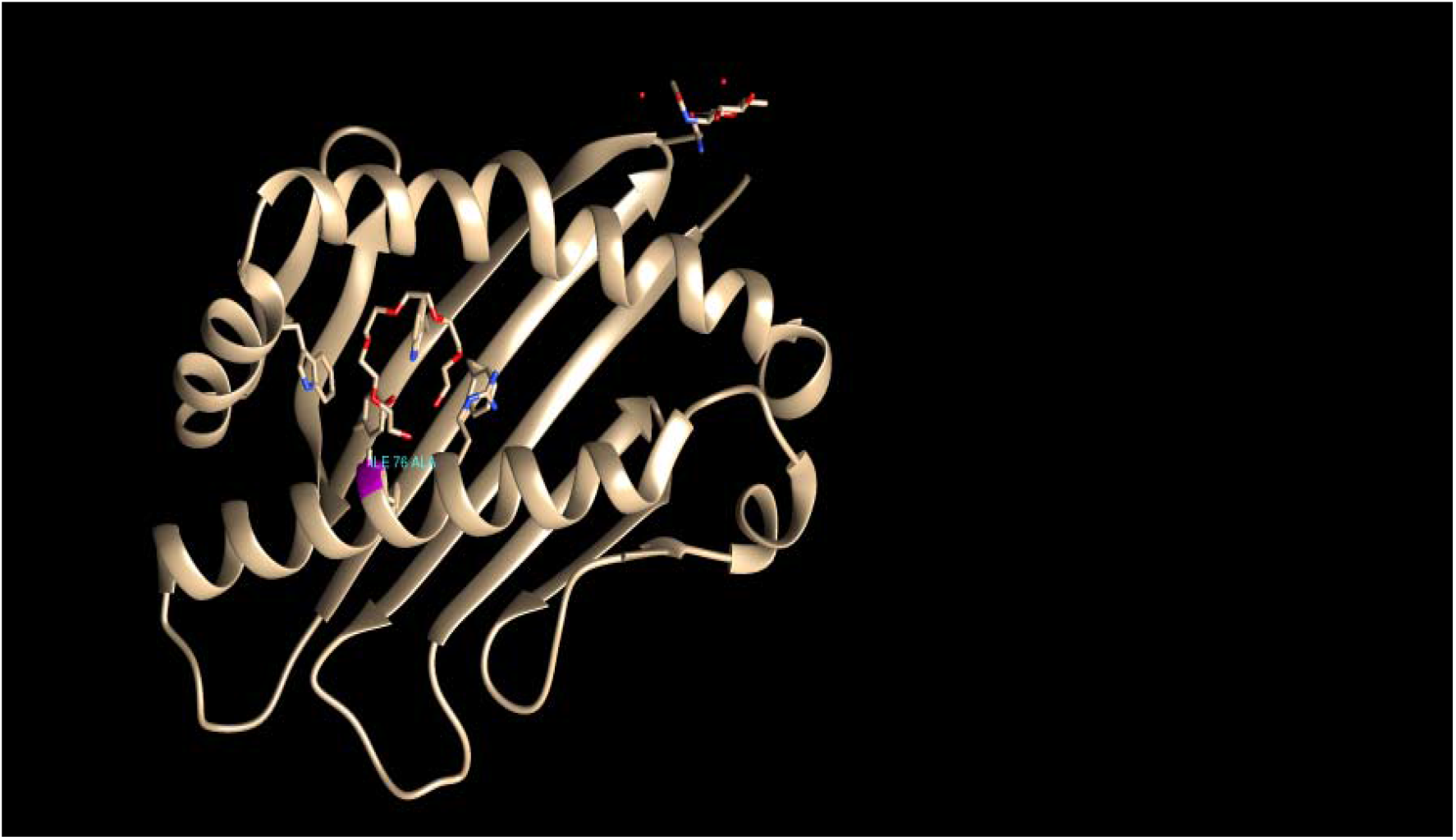
Zinc Alpha 2 Glycoprotein crystal structure obtained from protein data bank (1T7V) showing the Isoleucine amino acid residue mutated to Alanine. Blue- Tryptophan, Red -Phenylalanine, Yellow- Arginine, Magenta- Isoleucine.

**Figure 6.**
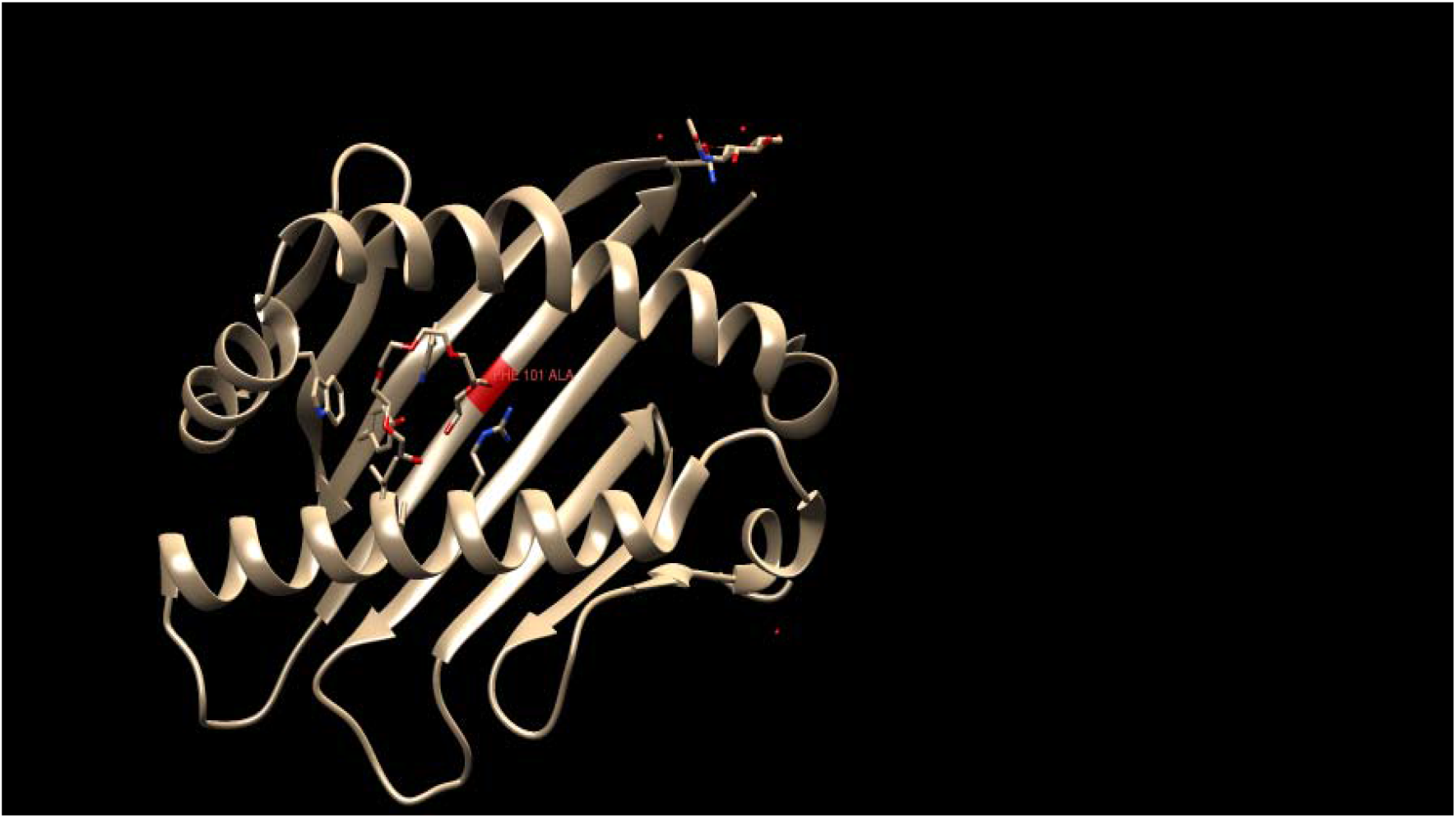
Zinc Alpha 2 Glycoprotein crystal structure obtained from protein data bank (1T7V) showing the Phenylalanine amino acid residue mutated to Alanine. Blue- Tryptophan, Red -Phenylalanine, Yellow- Arginine, Magenta- Isoleucine.

### Confirmation of ZAG pQE60 plasmid construction

pQE60 with ZAG expressing part were constructed with ZAG with no translational modifications in BL21 *E. coli* cells **(figure7)**. The BL21 cells are chosen for transformation as these cells are engineered for protein expression but are deficient in proteases so the digestion of recombinant proteins is lower than many other strains. BL21 *E. coli* competent cells are deficient in cytoplasmic protease Lon which is nuclear encoded mitochondrial ATP dependent serine peptidase and an outer membrane protease OmpT which is aspartyl protease. These mutations result in decreased protein breakdown and allows for increased protein accumulation within the transformed cells. Therefore, it produces a greater expression level of recombinant proteins compared to many other cells [23]. Each plasmid also contains an ampicillin resistance gene which allows the transformed cells to grow in presence of ampicillin whilst the wild type (WT) cells will not grow. The colonies grown on the plates will be ampicillin resistant. The plasmid also has Lac operon region which is promoter and activate in the presence of Lactose and absence of glucose in ZAG m-RNA expression. Colonies had grown on the plates of pQE60 BL21 cells. The transformation was successful and BL21 cells had taken up the plasmid with ampicillin resistant gene and had grown in presence of ampicillin. Control showed no growth while mutants arginine and tryptophan showed colonies. Restriction digest assays were then performed to ensure the bacteria growing were expressing the correct plasmids and the restriction enzymes were cutting at the correct size. pQE60 digest experiments resulted in single cuts with single digests using Bam HI, Eco RI, Bgl III and Hind III whilst a double digest of Bam HI + Bgl III resulted in two cuts producing two bands and Eco RI + Hind III only resulting in a single cut. The simulated version of the restriction digest analysis was being carried out using Snapgene to validate our own restriction digest gel. The lab experiments gave the same results as the virtual digest gels, supporting the hypothesis that successful transformation of the noted plasmids was achieved **(figure 8,9).**

**Figure 7.**
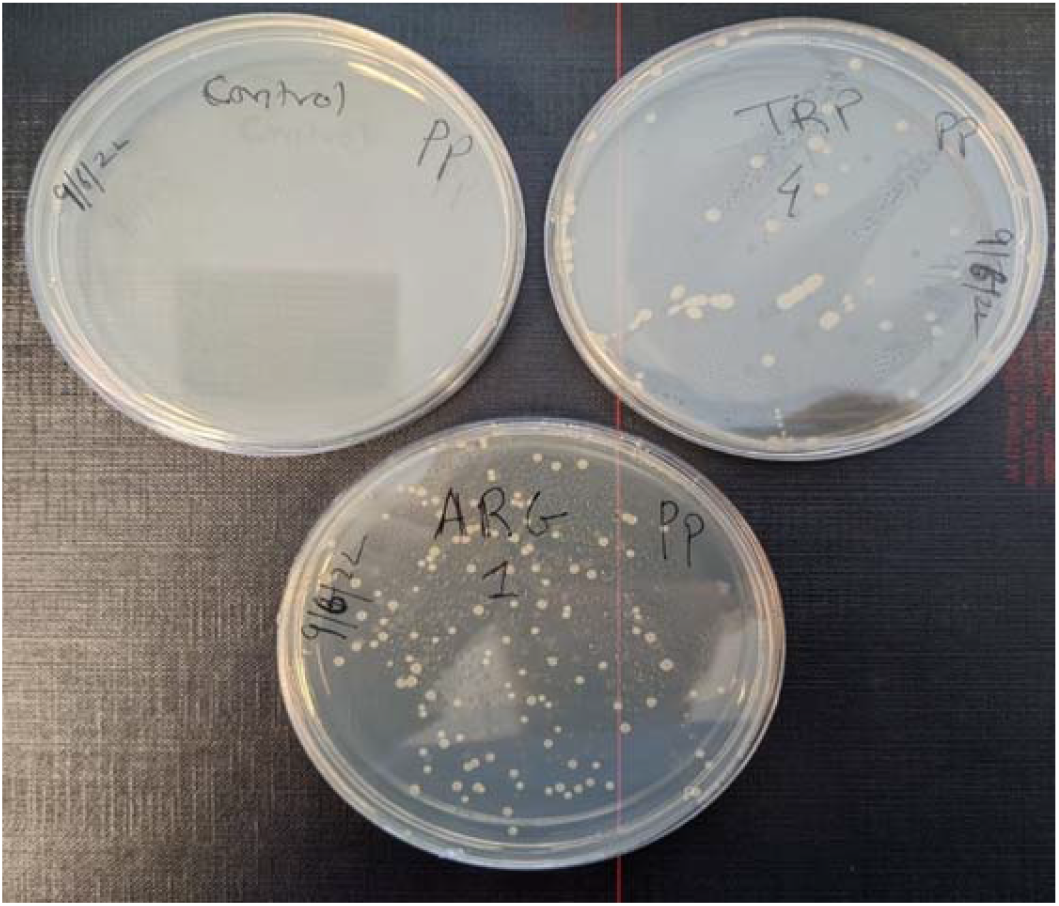
Plate map of mutants of Arginine and Tryptophan with colonies depicting growth of cells with ZAG region in it. Control plate showed no growth of cells. BL21 cells showed significant levels of ZAG after keeping overnight.

**Figure 8.**
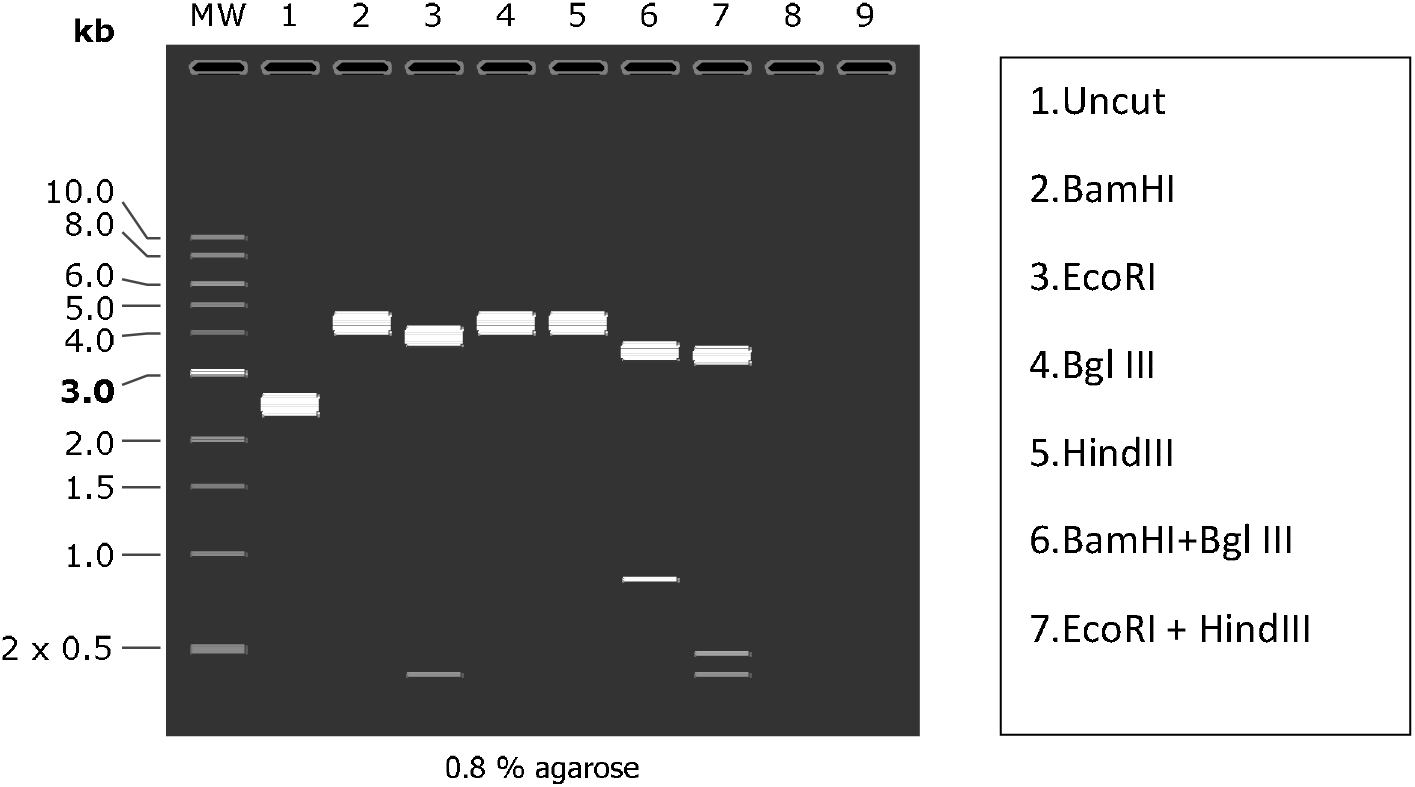
Virtual restriction digest gel run for PQT60 samples on 0.8% Agarose gel created on Snapgene.

**Figure 9.**
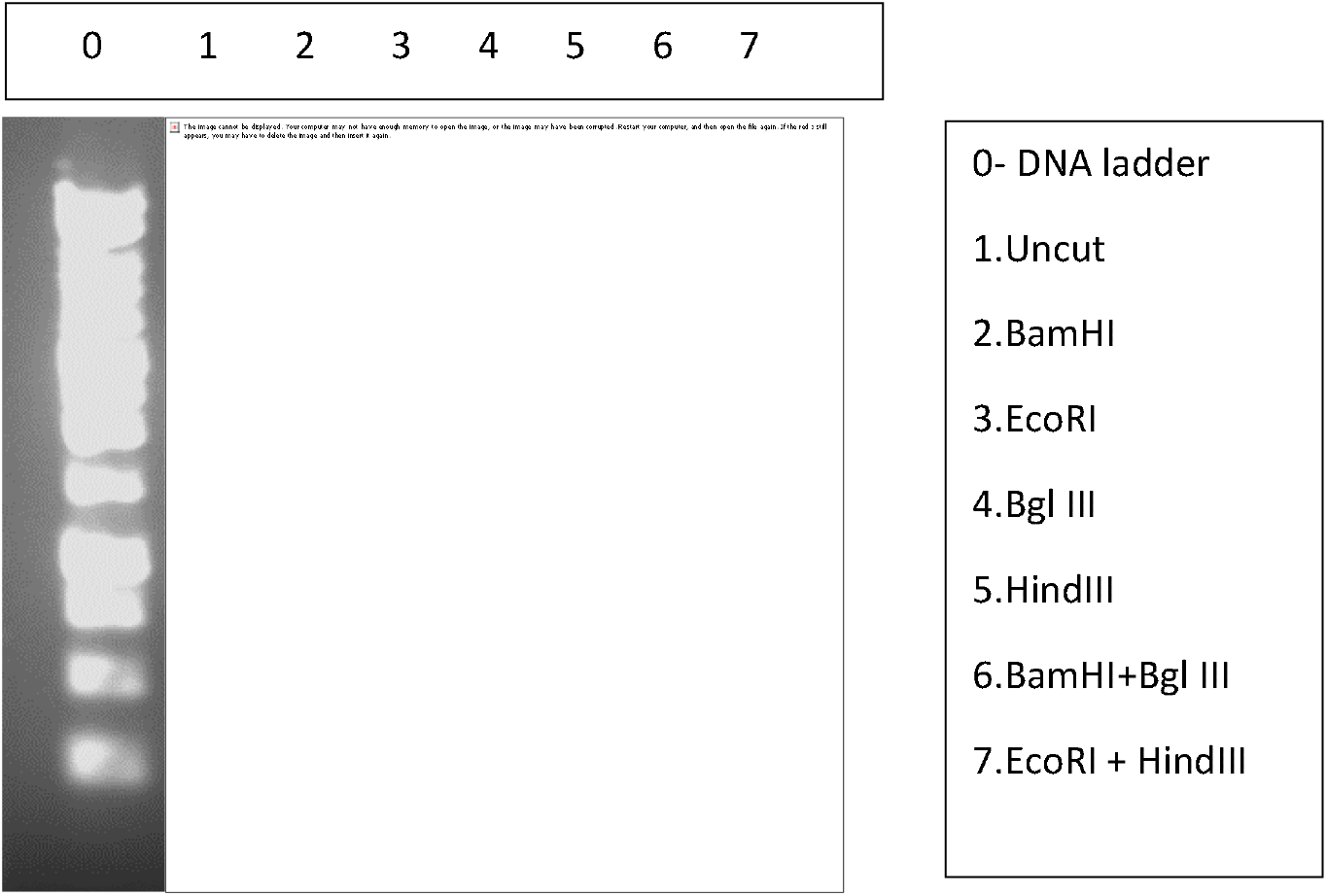
Restriction digest gel of DNA on agarose gel of pQT60 plasmid. BL21 competent cells were transformed with pQE60 plasmids, grown to OD600 and miniprep experiments were performed to obtain the DNA from both transformed BL21 cells. pQE60 plasmid underwent single digests with Bam HI, Eco RI, Bgl III and Hind III and double digests of Bam HI + Bgl III and Eco RI + Hind III as shown using the key on the right of the figure. Plasmids digests were analyzed on an agarose gel under U. V light. Faster migrating bands stain less intensely and are less clear on this image than is the case on illumination.

### Breaking the inclusion bodies to release ZAG

When recombinant proteins are expressed in high quantities in bacteria, this often results in insoluble and misfolded protein being packaged in inclusion bodies. This is due to the foreign nature of the protein, with cysteine residues forming disulphide bonds which the bacteria systems do not support. The extraction of ZAG from such inclusion bodies after undergoing lysis and detergent steps is crucial step as it facilitates purification of ZAG. The cells suspended in lysis buffer before the addition of lysozyme is important or else the unlysed cells will be spun down with inclusion bodies and will be contaminated with cellular proteins. Supernatants after each subsequent triton wash were collected and the final pellet after the triton wash steps expressed a large yield of ZAG **(figure 10,11)**.

**Figure 10.**
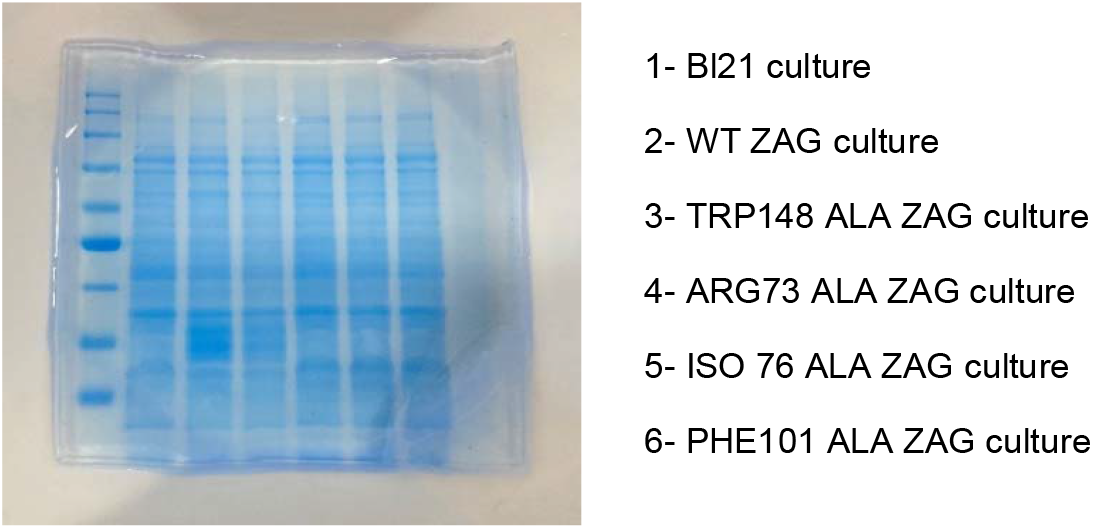
Validation of the transformed ZAG cultures with the help of Bl21 E.coli cells. The sample loaded was 10μL with MW marker of 3μL added to the respective lanes.The transformed cultures show a lot of hazy blue stain on the blot showing the cultures contain a lot of cell material and protein and debris.

**Figure 11.**
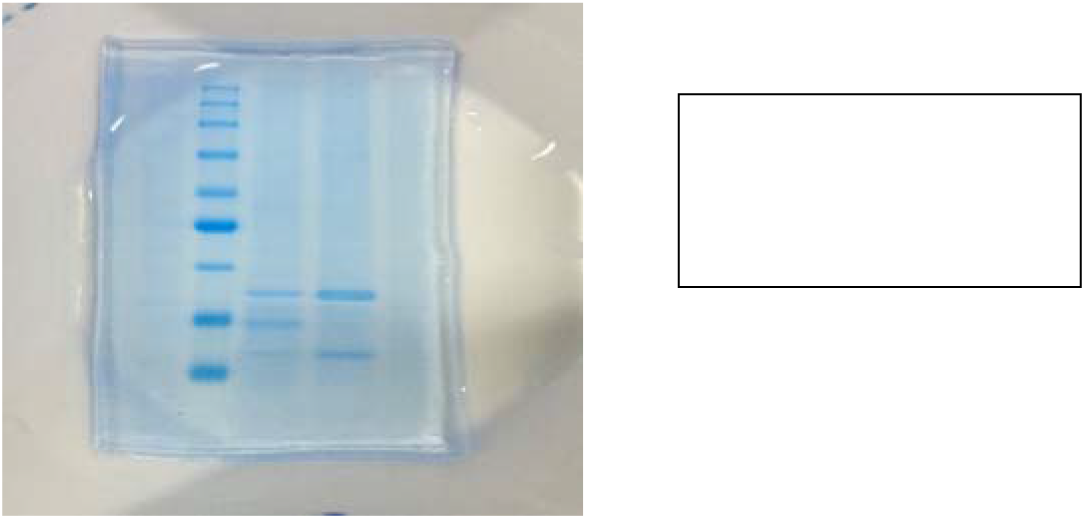
The gel is stained with instant blue stain for visualizing the proteins. This blot shows the protein is squeezed out fom the inclusion bodies by breaking them. Tryptophan looks double the size of arginine which shows unequal sign of loading. The sample loaded was 10μL with MW marker of 3μL added to the respective lanes.

### Denaturation and Refolding of protein and purification of the protein

After the exclusion of the protein from the inclusion bodies by the protein purification process, the protein undergoes denaturation and refolding. The ZAG protein produced in E. coli cells is misfolded and as a result has to be refolded for it to be biologically active. After the Triton washes it shows ZAG protein in the pellet. Further after denaturation and refolding step both cell lysates and supernatant are recovered. Denaturation by Guanidium Hydrochloride is due to interaction with polar parts of the protein. The small hard black pellet expressed a small concentration of ZAG in comparison to the pellet before the denaturing and refolding steps. The supernatant however, resulted in a significantly increased ZAG concentration in comparison to the pellet. BL21 expressing ZAG was used a positive control. These results imply that ZAG has been released from the inclusion bodies and the next procedure would now be to isolate the ZAG protein to produce a homologous sample for experiments. The purification was due to the vivaspin experiment used. The refolded ZAG contained in solution is spun, the ZAG stays in the top compartment of the tube and everything else passes through the filter into the bottom of the tube. The gel stained with instant blue validates the before and after protein purification steps **(figure 12).**

**Figure 12.**
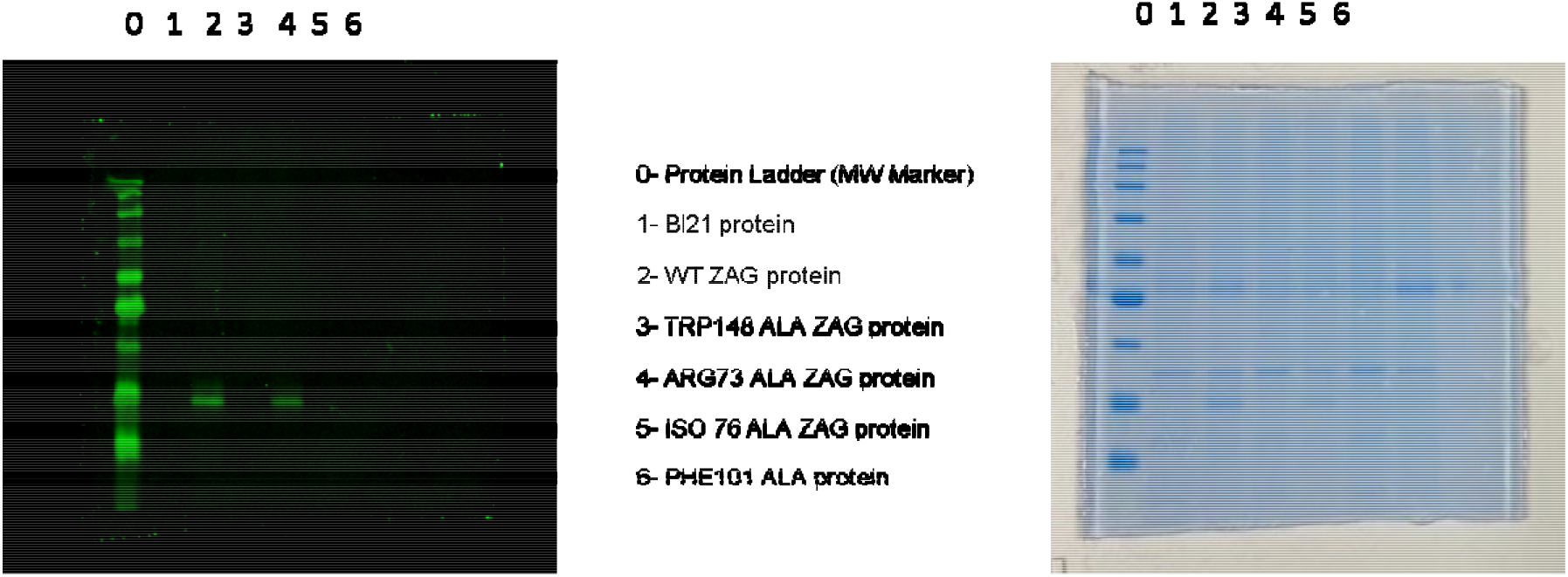
Western blotting of the protein samples of wild type as well as mutants. The gel on the right is stained with instant blue to visualize proteins. This blot shows the presence of ZAG in both wild type and arginine. The samples taken above are purified but not gone through a vivaspin process. The blue stain shows protein in the lanes of arginine-alanine mutant and wild type. The blank in the lanes of the blot shows there was no loading issues. The sample loaded was 10μL with MW marker of 3 μL added to the respective lanes.

### Lipolysis assay

The lipolytic effect of ZAG in developed adipocytes is observed in comparison with positive control isoproterenol. The 3T3-L1 adipocytes are treated with reagents for 10 days as mentioned in methods section. The time period plays a major factor in this experiment. The treatment is done for 1h with WT ZAG (1.5mg/ml), TRP-ALA ZAG (0.5mg/mL), ARG-ALA ZAG (3.2mg/mL). As per the results, the graphs depict that tryptophan mutation increases the lipolysis compared to wild type and arginine mutant **(figure 13)**.

**Figure 13.**
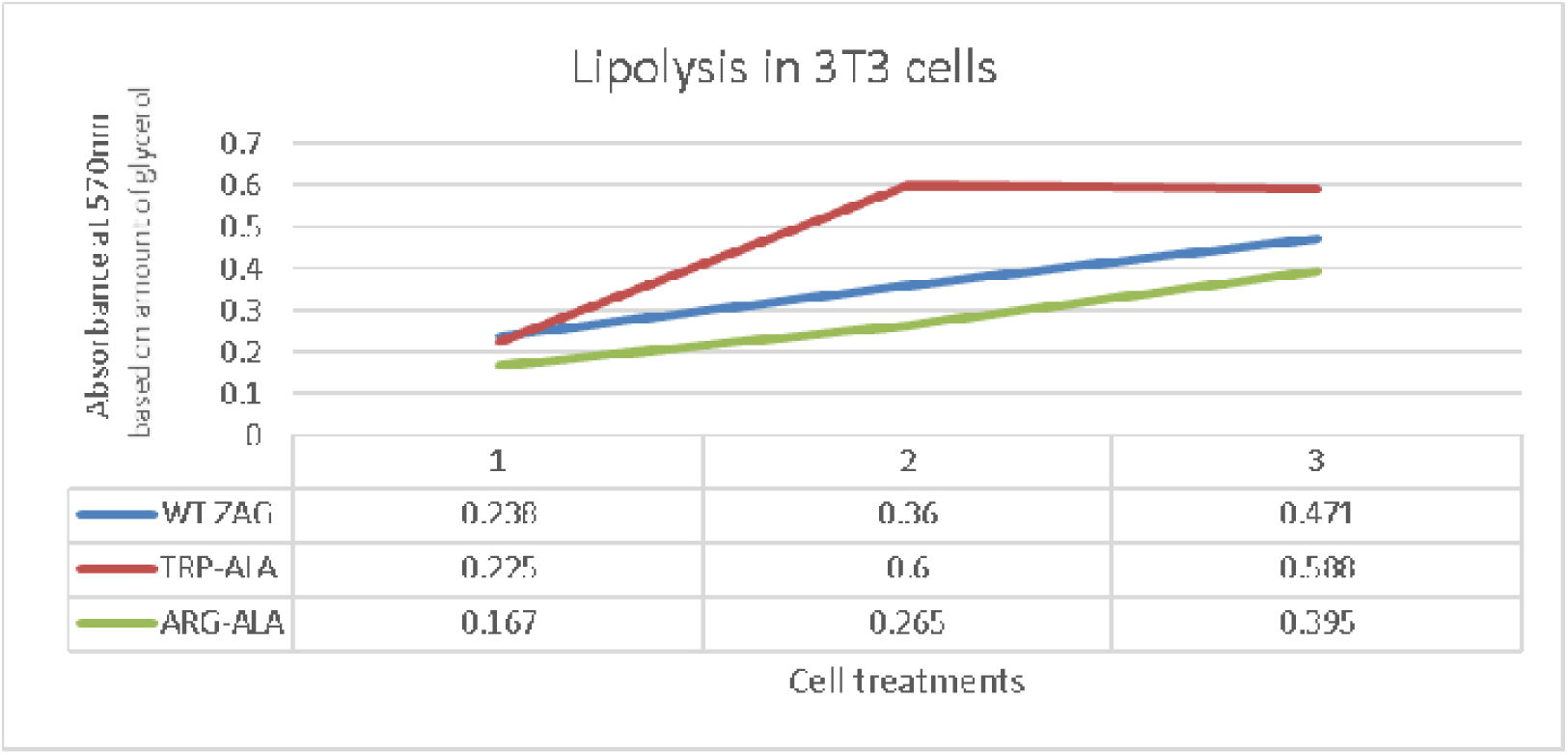
Line graph depicts the lipolysis in 3T3 adipocytes with a help of a colorimetric assay at absorbance of 570nm. Tryptopha mutant has shown a great increase in lipolysis with increase in concentration of the treatment.

### Oil red O staining

This staining technique is used for quantitative differentiation of 3T3-L1 adipocytes. The adipocytes are treated in the same way as for lipolysis experiment. On the day 10, the adipocytes are treated with concentrations of wild type ZAG 18.4μL, TRP-ALA ZAG 120μL, ARG-ALA ZAG 40μL. The treatment period in this scenario is 24 h unlike 1 h for lipolysis. The pictures after eluting the oil red stain with isopropanol were taken (**figure 14**). The red spots appear on the cells are triglycerides lipid droplets in the adipocytes. The spectrophotometric assay has been used to calibrate the amount of lipid droplets present in each category. These plate with absorbance mapped suggests there is no change in lipid content after 24 h **(figure15).**

**Figure 14.**
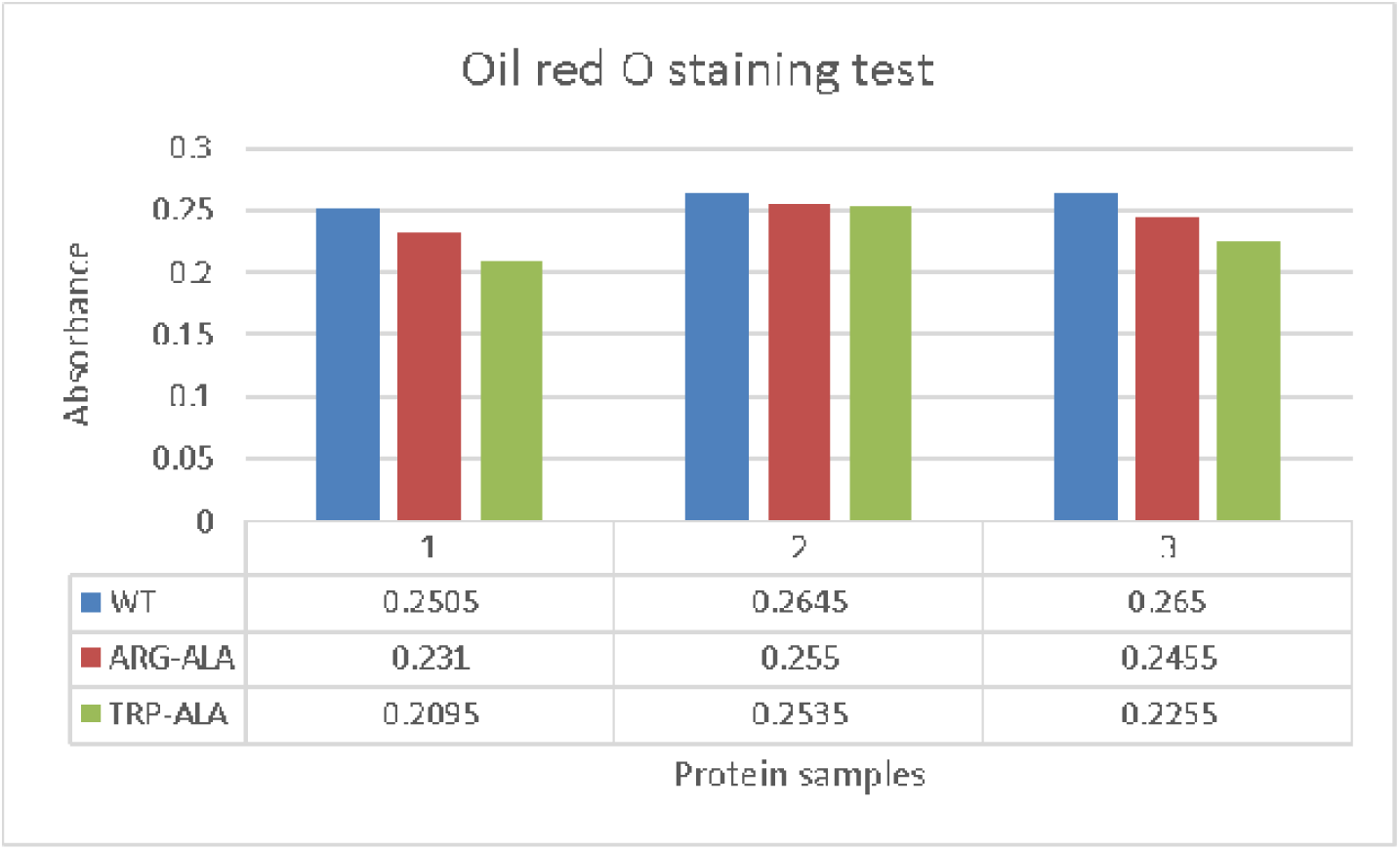
The graph above suggests the absorbance for Oil red O experiment for wild type and tryptophan and arginine mutant. The graph depicts no considerable change in lipolysis for each mutants and wild type.

**Figure 15.**
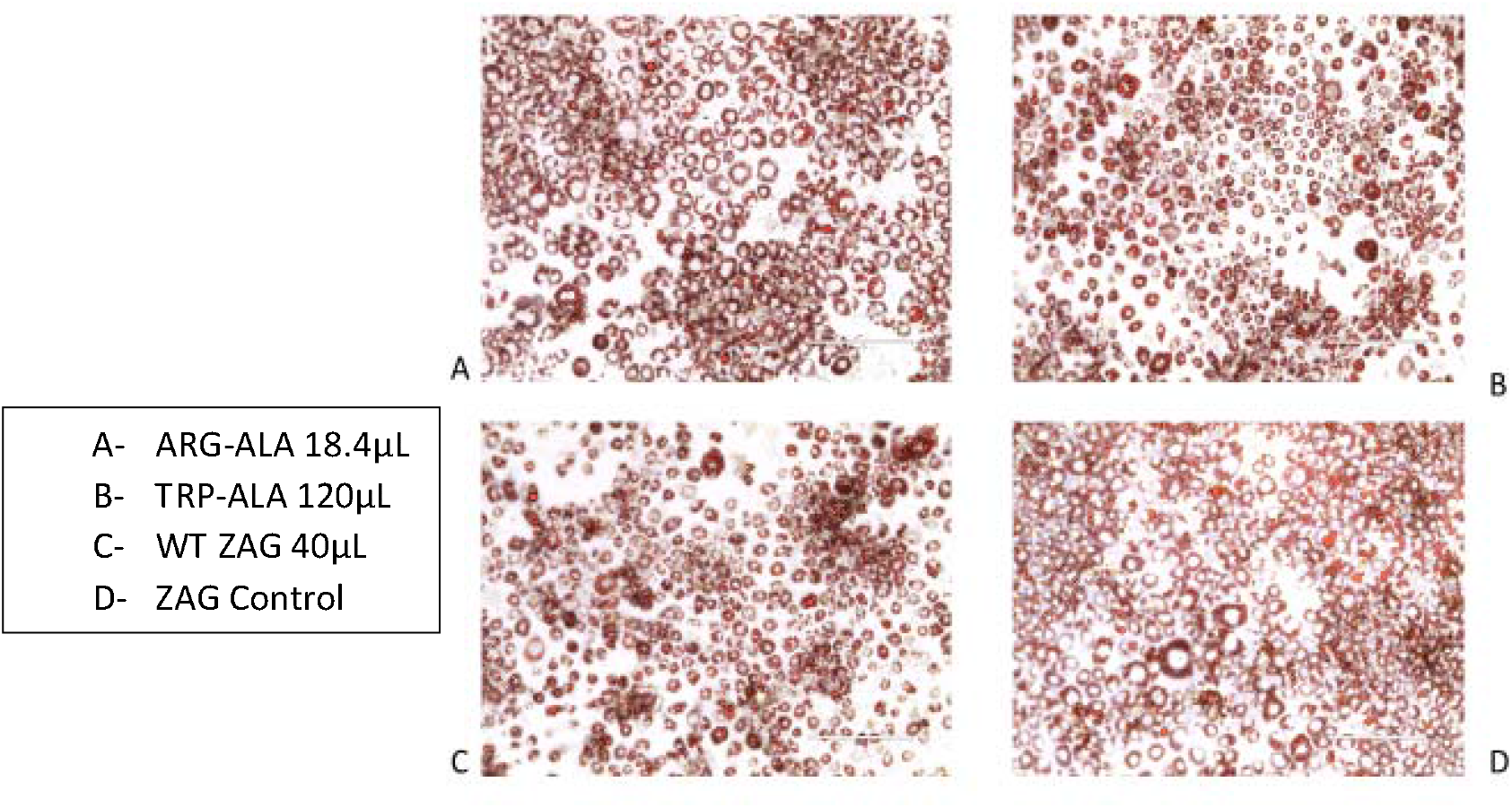
Oil red O staining pictures of the treated 3T3 adipocytes of wild type, control, tryptophan and arginine after day 10 for 24 h ZAG treatment. The triglyceride droplets are observed and stained after eluting red stain.

## Discussion

ZAG is regarded as the protein with many unknown functions. Adipose tissue secretes adipokines such as Zinc Alpha 2 Glycoprotein (ZAG). This protein shares the structural homology with MHC class I peptides. It is proposed that fatty acid ligands bind to the ZAG groove. ZAG has extremely comparable biological activity to LMF, and biochemical data suggests that both compounds promote lipolysis in vitro via a cyclic AMP-mediated mechanism via contact with a β3-adrenoreceptor) **(figure2,3,4,5,6)** [24]. β3-adrenoreceptor agonists induces adipolysis through classical cyclic AMP and protein kinase A pathway, and through the Extracellular Regulated Kinase (ERK) pathway, which sums up to 15% to 25% of lipolysis [25]. Just like ZAG, LMF also increased the glycerol levels and fatty acid levels after lipolysis. Protons and numerous anions are transported across the inner mitochondrial membrane by the ability of UCP-1. It promotes thermogenesis and aids in the uncoupling of respiration by converting ADP to ATP. Upregulation of UCP-1 is observed in BAT but not in WAT by ZAG which is mediated through a β3-adrenoreceptor [26]. Whereas tumour necrosis factor (TNF-α) which is also termed as cachectin downregulates UCP-1 in BAT by phosphorylation of peroxisome-proliferator activated receptor γ (PPARγ) via ERK pathway [27]. HSL is upregulated in cachectic patients which is similar to ZAG levels in obese patients. The m-RNA expression of HSL and ATGL is controlled by ERK pathway [28]. ZAG is considered as the driver in loss of adiposity. It is upregulated in cancer related to adipose tissue. This increase in results in weight loss which cannot be managed by diet and is observed majorly in the fourth stage of cancer. It is regarded as a side effect which cannot be cured.

As the experiments conducted above depicts the point to study the effect of ZAG on lipolysis and the role of mutants in the mechanism of cachexia. The difficulty in purifying this protein is the involvement of the inclusion bodies. As mentioned, protein production in bacteria can result in protein being misfolded and packaged in these vesicles. These inclusion bodies have to be taken up from the cells and the protein should be purified to isolate the desired protein. Each subsequent step in purifying ZAG has been optimized on a small scale to identify the most suitable method in producing ZAG from bacteria. Many papers have mentioned as DH-5α cells as the best human cells for transformation. We have used BL-21 E. coli cells as the it gives better output of ZAG production than DH-5α cells [23]. The optimum temperature of 37°C also helps in giving a better yield of ZAG. PQE60 plasmid has a lac operon DNA region inserted to help with protein production. When allolactose is present, it removes a repressor from the operator and initiates gene expression via the lac operon. The lac operon has been inserted next to the ZAG expressing region to upregulate ZAG production in the presence of allolactose **(figure 7,8,9**).

ZAG is carefully removed by performing each lysis step. After protein purification, in each triton wash step a small amount of ZAG is lost. The process of washing was tweaked as it is changed to 5 times rather than doing 3 washes. In previous methods we have used 6M Guanidium Hydrochloride for denaturation and removal of desired proteins [29] [30].The large band on the blot suggests that the Guanidium is working in this regard but it does not mix well with the SDS-PAGE buffer. When performing western blotting, the sample turns solid and need repeated heating at 95° C as it enters the well during loading and is the reason for the smudged effect. The denaturation step is then carried out using 8M urea as it is observed to give clearer and cleaner blots. The ZAG protein along with all the mutants have undergone vivaspin in the vivaspin columns. The vivaspin is carried out at maximum speed and the supernatant is checked for protein concentrations. The ZAG mutants such as tryptophan and phenylalanine when purified formed not so harder pellets unlike arginine and isoleucine. Thus, it can be hypothesized that mutation with any amino acid may or may not alter the function of protein and the refolding process of the protein [31, 32]. The washing and denaturation step can be clearly pointed out. The cultures when blotted and after the protein purification step can show clearer blots in arginine and tryptophan than the transformation blot **(figure 10,11).** The blot which is visualized using green rabbit antibody shows presence of ZAG in wild type and Arg mutant. The samples for this blotting have gone through the same protein purification process with a slight change of urea for denaturation. These samples have not gone through vivaspin thus there is marked low concentration of ZAG in the samples. These experiments prove the point of need for vivaspin and change in reagent to urea causes a lot of change **(figure 12).**

Lipolysis is the hydrolysis of triglycerides into free fatty acids and glycerol. Adipose TG lipase and HSL, as well as monoglyceride lipase, are involved in this process. The use of isoproterenol is best as it is a catecholamine that activates β-adrenergic receptors. This leads to activation and converts ATP to c-AMP. Cyclic -AMP then activates the hydrolysis of triglycerides by HSL [33]. As per the hypothesis, ZAG treatments have also been found to increase the levels of adipose TG lipase and HSL, which contribute to lipolysis [34][35]. Overexpression of ZAG also reduces the mRNA levels of fatty-acid synthase (FAS), acetyl-CoA carboxylase, and acyl-coenzyme A while increasing the mRNA level of hormone-sensitive lipase and thus these cumulative effects leads to suppression of lipogenesis and lipolysis enhancement. The graph suggests that tryptophan mutant shows increased lipolysis after 1 h than wild type and arginine mutant **(figure 13).**

The Oil red O dye is selective as it only stains the triglycerides and cholesterol oleate [36]. There is no change in the lipolysis content after 24 h. This suggests that time plays a major role in expression of ZAG. The 24 h showed no lipolysis whereas the 1 h showed considerable change. 24 h of ZAG treatment can be considered as prolonged term than short period of 1h for expression of glycoprotein **(figure 14,15).**

The future plan is chalked out for probing ZAG and its effect on adipose tissue. The western blots will be carried out to find out the effect of ZAG on the enzymes such as MAP kinase, Hormone Sensitive Lipase, Fatty Acid Synthase. The upregulation and downregulation of the enzymes clearly show that ZAG wild type and mutants have a distinct effect on the lipolysis. The following work would be to look at the cell signaling and observe the differences between wild type and mutants. Hence the current study demonstrated that the Tryptophan-Alanine mutant shows increased lipolysis after 1 h. The wild type and Arginine-Alanine mutant does not show a comparable rate of lipolysis. The time period of 1 h can also play a major role in showing the effect. As discussed, cachexia results in the patients becoming very weak and the resources of chemotherapeutics are limited due to the frail state of the patient and the toxic nature of the treatments. The design of a therapeutic would give patients options of treatments brought about by attenuating the weight loss. With this, it would offer a better prognosis for patients and provide them with a greater quality of life.

